# Non-destructive extraction of DNA from preserved tissues in medical collections

**DOI:** 10.1101/2021.02.17.431618

**Authors:** Enrique Rayo, Giada Ferrari, Judith Neukamm, Gülfirde Akgül, Abagail M. Breidenstein, Martyn Cooke, Carina Phillips, Abigail S. Bouwman, Frank J. Rühli, Verena J. Schuenemann

## Abstract

Museum and medically fixed material are valuable samples for the study of historical soft tissues and represent a pathogen-specific source for retrospective molecular investigations. However, current methods for the molecular analysis are inherently destructive, posing a dilemma between performing a study with the available technology thus damaging the sample - or conserving the material for future investigations. Here we present an unprecedented non-destructive alternative that facilitates the genetic analysis of fixed wet tissues while avoiding tissue damage. We extracted DNA from the fixed tissues as well as their embedding fixative solution, to quantify the DNA that was transferred to the liquid component. Our results prove that human ancient DNA can be retrieved from the fixative material of stored medical specimens and provide new options for the sampling of valuable curated collections.

**Method summary:** We compared the metagenomic content of historical tissues and their embedding liquid to retrieve DNA from the host and specified pathogens based on the diagnosis of the sample. We applied ancient DNA research techniques, including in-solution hybridization capture with DNA baits for human mitochondrial DNA, *Mycobacterium tuberculosis, Mycobacterium leprae*, and *Treponema pallidum*.

## Main text

Fixed wet tissues from museums and anatomical collections offer an extensive, mostly pathogen-specific, and precisely dated archive for retrospective molecular investigations. Although most collections are accessible for collaborative, scientific inquiry, in the case of very unique or valuable material, destructive methods such as sampling for molecular analysis are not permitted. For this reason, developing non-invasive sampling methods for specimens in museums or medical collections has been a central objective of the paleogenetics field for decades (Cobb 2002; Thomsen et al. 2009; Rohland, Siedel, and Hofreiter 2004), but even in those cases a minimal intrusion and retrieval of organic material is necessary. Traditional extraction techniques (Rohland and Hofreiter 2007) rely on the binding of DNA strands to inorganic structures, like bone or teeth, or the presence of enough preserved genetic material within soft tissues. The retrieval of DNA is then achieved by homogenizing and chemically dissolving the tissue. However, DNA from organisms has been reported to survive freely in an array of different ecosystems, such as soil, freshwater, or saltwater, and became known as environmental DNA (eDNA) (Ficetola et al. 2008; Taberlet et al. 2012). This raises the question of whether the same leaching principle is observable with specimens preserved in embedding liquid for a long period of time, such as the case for medical and archival collections, where tissues of interest are preserved in the same ethanol or formalin preservative solution for decades or even centuries. If DNA leaches into the fixative liquid, it would represent a less invasive substrate to sample to obtain human and pathogen DNA for molecular analyses than the tissue itself. The molecular detection of leached DNA from a preserved specimen has been tested previously, but only with traditional PCR methods and very recent material - 7 ∼ 10 years old (Shokralla, Singer, and Hajibabaei 2010). To our knowledge, no other study has assessed the feasibility of characterizing DNA utilizing the embedding liquid of historical collections.

Here we present results from an exploratory study on fixed specimens from the Museum collections at the Royal College of Surgeons (RCS) in London (United Kingdom). These collections contain approximately 3,500 18th century preparations from the original collection of surgeon and anatomist John Hunter, and 7500 other preserved tissue preparations ranging from the 19^th^ century to the present day. We collected tissue and fixative samples from ten specimens (14 in total with some replicates subsampled) dated between 1760 and 1886 CE. The selected specimens were diagnosed with tuberculosis, leprosy, syphilis, cancer, and skin conditions (Table SI.1). These historic samples represent good candidates for a retrospective genomic analysis of these disease-causing pathogens, namely members of the *Mycobacterium tuberculosis* complex (MTBC), *Mycobacterium leprae*, and *Treponema pallidum* subsp. *pallidum*. With metagenomic shotgun sequencing data obtained from tissues as a baseline, we assessed the retrieval of human and candidate pathogen DNA from the corresponding fixatives, using both a shotgun sequencing approach and hybridization capture of the human mitochondrial (mtDNA) and bacterial pathogen DNA (overview in Fig.1) (SI Notes 1&2).

**Figure 1:**
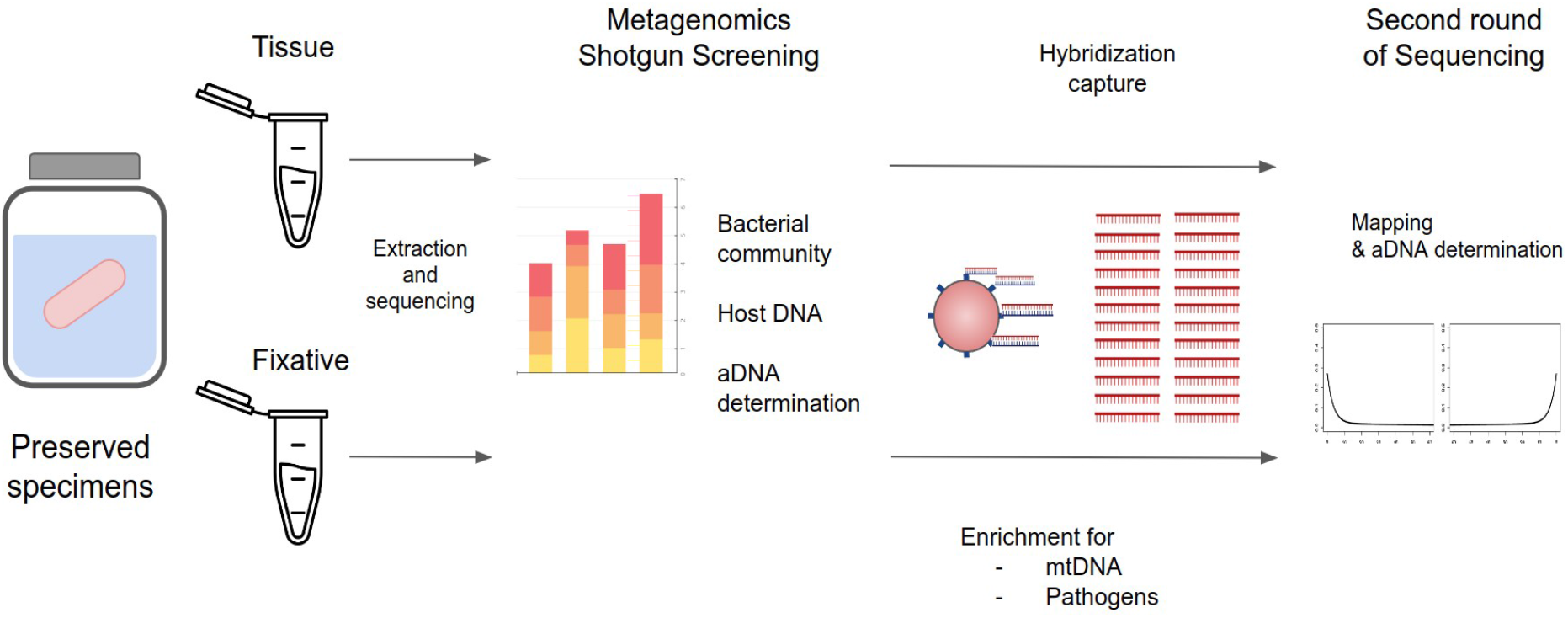
Schematic representation of the workflow of the project.

For the first assessment of ancient DNA preservation in the samples, we screened the shotgun sequencing output for reads that could be mapped against the human mtDNA. Nine out of 14 tissue replicates showed detectable traces of mtDNA, eight with a 5X coverage above 98% and one above 24% (Table SI.5). These replicates demonstrate the characteristic misincorporation patterns for ancient DNA (elevated C to T substitutions at 5′-ends and elevated G to A substitutions at 3′-ends) in frequencies ranging from 10-20% damage (Neukamm, Peltzer, and Nieselt 2020). On the other hand, only the liquid sample HA4.1 showed DNA that could be considered as ancient (94.59% 5X coverage of mitogenome, 16% damage). The enrichment strategy (Maricic, Whitten, and Pääbo 2010) (SI Note 2) was powerful enough to increase the endogenous content of all the tissue samples (Table SI.6), with fold increments from 14 to 275, and also for eight of the ten liquid samples (Table SI.7), not including HA4.1. This sample had a remarkable increase with a final output of more than three million mapped ancient mtDNA reads, a 537-fold increase in the endogenous content (100% coverage), and retaining damage of 10% (Fig.2A).

Only seven tissue samples and one liquid sample (HA4.1) had sufficient reads and coverage for mitochondrial haplotype assignment and contamination estimation (Table SI.8). The HA4.1 tissue sample was characterized as the Eastern European haplogroup H2a2, and the corresponding liquid sample was assigned to the same haplogroup but to a higher definition due to the high number of reads retrieved (H2a2 subgroup a1), confirming the transfer of mtDNA from the tissue to the embedding fixative material. An estimated 0.01% of reads from HA4.1 were identified as contaminated by haplogroup K1 (Renaud et al. 2015). This could have happened due to handling - either during tissue preparation or further curation treatments (e.g., refilling the vases), but it is still a very low percentage of contaminant reads. For the rest of the tissue samples, the haplogroups identified were also typically European, which was not a surprise since the samples were expected to be of European origin (Table SI.8) (Blair 2013; Weissensteiner et al. 2016).

Bacterial detection for shotgun screening, on the other hand, was less satisfactory. The metagenomic analysis showed certain sample-dependent variation in the number of mapped bacterial reads, with a general higher abundance of bacterial reads in the tissue (from 300,000 to 1,200,000) as compared to the ethanol (ranging from 50,000 to 650,000) (Fig.2B). We estimated richness by calculating the bacterial alpha diversity using the observed and Shannon indexes (McMurdie and Holmes 2013). Tissue samples consistently scored higher for both indexes (Fig. SI.1).

At the Phylum level, the tissue samples are heavily dominated by Proteobacteria (from 55% to 99% of total reads). Liquid samples are also dominated by Proteobacteria (20% ∼ 78%), but with Firmicutes (53% in some samples) and Actinobacteria (12% ∼ 32%) in higher relative abundance (Fig.2C). At the Family level, some tissue samples were heavily dominated by members of Enterobacteriaceae (75% ∼ 95%), while others were higher for others such as Pseudomonadaceae (20%), Comamonadaceae (7% ∼ 20%), and Burkholderiaceae (5% ∼ 15%). The fixative samples had a more homogeneous abundance distribution of the main families, with Burkholderiaceae (10% ∼ 27%), Clostridiaceae (5% ∼ 55%), and Micromonosporaceae (5% ∼ 28%) present among all samples. A noticeable presence of *Clostridium* in the liquid samples was detected, higher in the ethanol-based fixative (ranging from 4% up to 56%) but very low in the Kaiserling fixative HA4.1E (<1%). *Clostridium* was absent or barely detectable in the tissue samples. At the species level, there were not enough reads to be reliably mapped to the reference genome, and no taxa presented a damage pattern to be validated as ancient. These families mentioned above are widespread in most environments (Cousin 1999; Octavia and Lan 2014; Voronina et al. 2015), and members of *Clostridium* are nearly ubiquitous in nature and very resistant to heat, desiccation, and toxic chemicals (Figueiredo et al. 2020). This suggests that most of the bacteria in these samples were present due to environmental contamination. The only sample that shows a shared component of bacteria with its respective tissue sample is HA4.1, with a significant abundance of the families present in the tissue (Enterobacteriaceae, 22% and Methylobacteriaceae, 12%) (Fig. SI.2), and lacked the presence of *Clostridium* that was present in other fixative samples, while being the only fixative sample with similar bacteria as in the tissue samples, such as *Klebsiella*. When we applied SourceTracker2 (Knights et al. 2011) to identify the proportion of the liquid bacteria community explained by the tissue samples, HA4.1 had a higher presence of taxa from the tissues, contributing 40% of the bacteria present in the HA4.1 fixative; it is noteworthy that the main contributors in the liquid community were the blanks, contributing from 30% up to 70% of the bacterial community in the liquid. This shows the strong impact of trace bacterial DNA even from sterile reagents in low-mass DNA studies (Fig. 2D).

**Figure 2.**
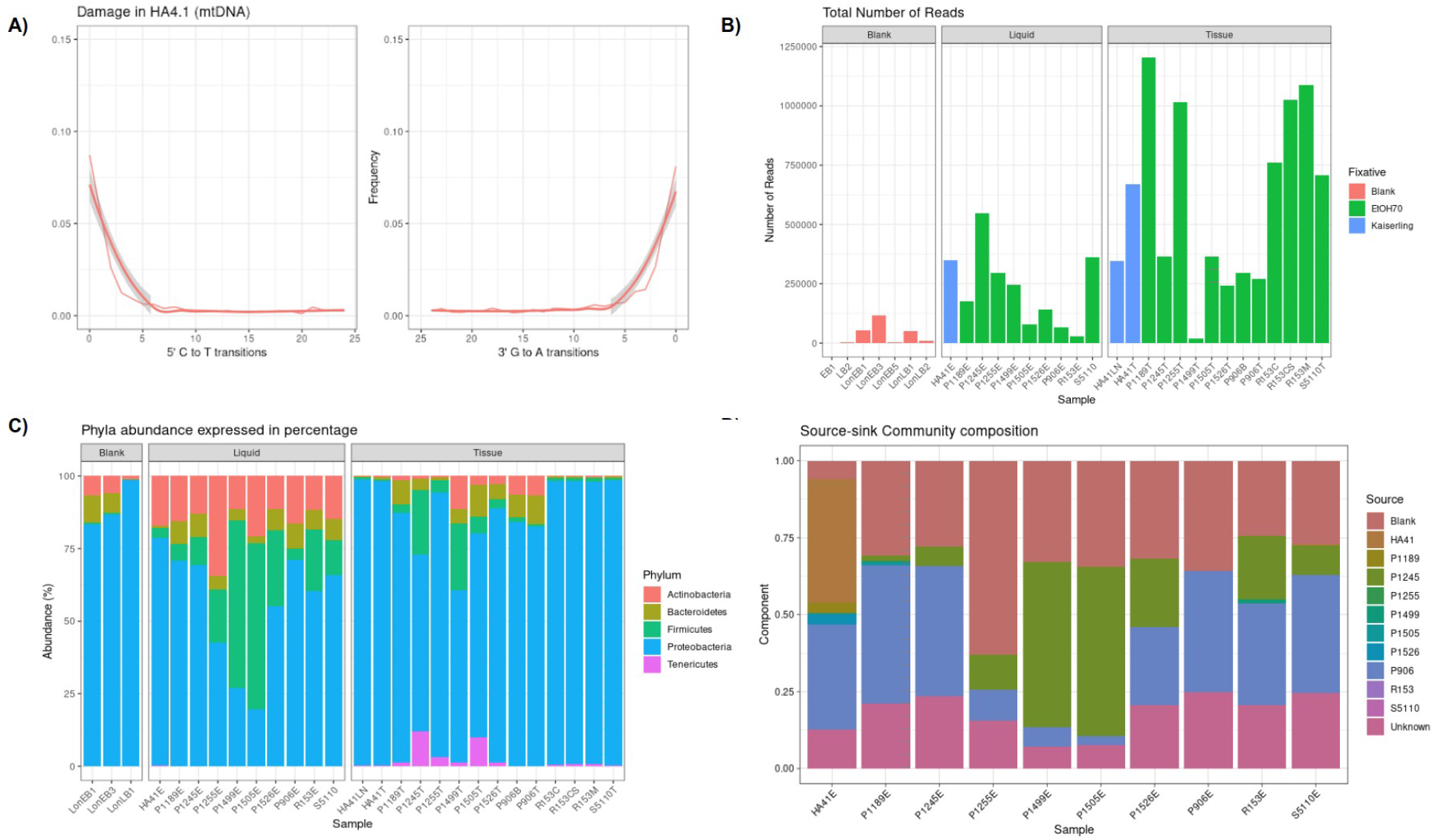
A) Damage pattern of the mtDNA reads from sample HA4.1 after hybridization capture. B) Total number of bacterial reads on each sample. C) Main bacterial phyla present in the samples, expressed in abundance. D) Bacterial community composition expressed in percentage component by SourceTracker2.

To test the viability of detecting pathogen DNA, we performed pathogen-specific hybridization captures for *M. tuberculosis, M. leprae*, and *T. pallidum* subsp. *pallidum* (SI Note 2). Only the *Mycobacterium tuberculosis* and *Mycobacterium leprae* enrichments were successful, while the *Treponema pallidum* protocol did not yield any usable output (Tables So. 9 and SI. 10). For the fixative material from HA4.1, only 790 reads were detected, with a damage pattern of 22% (0.082% endogenous DNA, 0.3 coverage), which did not provide enough material for a more in-depth analysis of the pathogen. Despite having the damage profile characteristic of ancient reads, with a cluster factor of 10.29 at 5% of a lane we considered that a deeper sequence would not provide better results to discern whether the reads are truly endogenous ancient *M. leprae* or correspond to environmental contamination presenting damage from other reasons that age (Bouwman et al. 2012). Given that the mtDNA enrichment worked for the tissues and one liquid sample, this means that the tissue samples themselves were not especially suited for the characterization of bacterial aDNA. Therefore, the testing of pathogen DNA leaching was not successful with this sample set.

Several limitations arise from this study: the success rate was low, with only one sample (HA4.1) presenting retrievable human reads from the fixative material, and that had a bacterial profile that is 40% similar to the one characterized in the HA4.1 tissue sample. As a small-scale study with limited sample size, this approach proved to be successful for extracting host aDNA, but the recovery of bacterial DNA - either commensal or pathogenic, was more complicated to achieve. Some potential solutions to improve the technique could be implemented for future studies, such as increasing the fixative volume extracted: as a screening, we extracted between 1-2 mL of liquid fixative, so we believe that a larger starting amount could increase the amount of DNA in suspension. In addition, as much detail as archives can provide, the maintenance and exchange of the fixative material were typically not recorded, where each exchange could potentially remove suspended DNA since samples with ‘older’ fixative liquid would theoretically have more aDNA. Also, it seems that the type of fixative could impact the transfer of DNA. When explaining this apparent success, we revisited the information detailing the fixative conditions (Table SI.1). Given that this sample was stored in Kaiserling solution instead of 70% ethanol may have contributed to the preferential preservation of free DNA in the liquid content. This fixative is generally an aqueous solution of formalin, potassium nitrate, and potassium acetate, although several variations were commonly found among collections (SI Note 4). In addition, it seems to be preventing the growth of environmental taxa better than the 70% ethanol solution, as shown by the lower presence of *Clostridium*. However, as is common with historical archival sources, the history of these samples is somewhat incomplete and it is possible that other liquid fixatives were added throughout the centuries (SI Note 4). Alternatively, it could be a simple matter of better preservation of this specific sample, independently of the type of fixative. Further selection and analysis of more samples preserved in Kaiserling solution would elucidate if this fixative procedure is a consistent and reproducible source for leached DNA.

In conclusion, here we present the potential use of liquid fixative materials from historical collections as a source for ancient DNA studies. We demonstrated that host aDNA can be retrieved from preserved samples without destroying the embedded tissue, opening a new array of possibilities for the molecular analysis of medical and historical collections.

## Author contributions

E.R. and G.F. conceived the presented idea. M.C. and C.P. provided samples and historical context. E.R., G.F., A.Br. and G.A. carried out the laboratory work. E.R. and G.F. wrote the manuscript with input from V.S, A.Br. and M.C.. J.N. and E.R. performed the metagenomic analysis and data intepretation. G.F., E.R. and A.Bo. obtained the funding for the project. V.S., F.R. and A.Bo.supervised the project. All authors provided critical feedback and helped shape the research, analysis and manuscript.

## Acknowledgments

We would like to thank the University of Zürich University Research Priority Program (URPP) and to Prof. Dr. Christian von Mering for the funding of this project, and thanks to the support of the Swiss Mäxi-Foundation in the completion of this project.

