## Supplementary Information for "Non-destructive extraction of DNA from preserved tissues in medical collections"

Enrique Rayo ^1^*, Giada Ferrari ^1,2^*, Judith Neukamm ^1,3^, Gülfirde Akgül ^1^, Abagail M. Breidenstein ^1^, Martyn Cooke^4^, Carina Phillips^4^, Abigail S. Bouwman^1^, Frank J. Rühli^1^, Verena J. Schuenemann^1^

1. Institute of Evolutionary Medicine, University of Zurich, Zurich, Switzerland;

2. Centre for Ecological and Evolutionary Synthesis (CEES), Department of Biosciences, University of Oslo, Oslo, Norway

3. Institute for Archaeological Sciences, University of Tübingen, Tübingen, Germany

4. Museums and Archives, The Royal College of Surgeons of England, London, United Kingdom

**Supplementary Note 1 - Laboratory workflow**

#### **Samples used in this study**

Samples were obtained from the Hunterian Museum collections at the Royal College of Surgeons in London (RCS) (UK) and are summarized in Table SI.1. We collected tissue and fixative samples from ten specimens (fourteen in total with some replicates subsampled) dated between 1760 and 1886 CE. The selected specimens were diagnosed with tuberculosis, leprosy, syphilis, cancer, or skin conditions. All samples are fully encoded and are older than 70 years (post-mortem). Therefore, they do not require additional approval under Swiss law (Swiss Federal Act on Research involving human beings. Human Research Act, HRA art.1 and art.36; RS 810.30) according to the responsible ethics committee (Kantonale Ethikkommission Zürich, Switzerland).

#### **Processing of tissue samples**

All DNA extractions and sequencing library preparations were performed in an ancient DNA clean laboratory (Knapp and Hofreiter 2010) following standard anti-contamination protocols (Cooper and Poinar 2000; Gilbert et al. 2005; Llamas et al. 2017) with parallel non-template controls.

The tissue was extracted following a protocol by Devault et al. with slight modifications (Devault et al. 2014). Approximately 100 mg of tissue were homogenized using a sterile scalpel blade and incubated at 95°C for 5 min in 800 µl of extraction buffer (25 mM Tris-HCl, 5 mM CaCl2, 25 mM sodium citrate, 2.5 mM ethylenediaminetetraacetic acid, 1% sodium dodecyl sulfate, 50 mM 1,4-dithiothreitol, 10 mM N-phenacylthiazolium bromide). After adding 80 µl of proteinase K (20 mg/mL), digestion was performed at 50°C and nutated for 24 h. A total of 1 mL was transferred to 10 ml of binding buffer (5M GuHCL, 40 % Isopropanol, 400 µl sodium acetate 3 M; UV light treated before use). The mixture was then transferred to a high pure extender assembly (Roche Applied Science, UV treated)  modified with a MinElute column (QiaGen), centrifuged and washed twice with 700 μl PE buffer before elution in 100 μl TET. Sequencing libraries were generated from 10 μl of extract or extraction blank following (Meyer and Kircher 2010; Kircher, Sawyer, and Meyer 2012) and sequenced on one lane of a HiSeq2500 (Illumina) with paired-end 125 bp reads and v4 chemistry by the Functional Genomics Center Zürich (Switzerland).

#### **Processing of fixative samples**

The fixative extraction was performed following a slight modification of the protocol by (Dabney et al. 2013). Approximately 1-2 mL of fixative liquid was sampled from the embedding liquid of each sample, and transferred without prior digestion to 10 mL of binding buffer (5M GuHCL, 40 % Isopropanol, 400 µl sodium acetate 3 M; UV treated before use). The mixture was then transferred to a high pure extender assembly (Roche Applied Science, UV treated)  modified with a MinElute column (QIAGEN), centrifuged and washed twice with 700 μl PE buffer before elution in 100 μl TET.

Sequencing libraries were generated following   (Meyer and Kircher 2010; Kircher, Sawyer, and Meyer 2012) with modifications. The re-amplification step was performed with 1 unit Herculase II DNA polymerase (Agilent), 5X Herculase II  reaction mix, 0.3 μM primers IS5 and IS6 (Meyer and Kircher 2010) and 4-7 μl library template with the following thermal profile: initial denaturation at 95◦C for 2 min, 10 to 25 cycles of denaturation at 95°C for 30 sec, annealing at 60°C for 30 sec and elongation at 72°C for 30 sec, followed by a final elongation at 72°C for 5 min. Libraries were purified with MinElute spin columns (QIAGEN) following the manufacturer’s instructions. Quantitative PCR (qPCR) and analysis on an Agilent 2200 TapeStation were used to assess the quality and concentration of the libraries. Libraries were pooled equally and sequenced on one lane of a HiSeq2500 (Illumina) with paired-end 125 bp reads and v4 chemistry by the Functional Genomics Center Zurich (Switzerland).

#### **In-solution capture with DNA baits**

After the shotgun screening, the samples were enriched by using a hybridization capture protocol with customs biotinylated baits for human mtDNA (Maricic, Whitten, and Pääbo 2010) and the pathogens *Mycobacterium leprae*, *Mycobacterium tuberculosis* (Schuenemann et al. 2013; Bos et al. 2015, 2014; Furtwängler et al. 2020), and *Treponema pallidum*. For the syphilis capture four genes from the tpr gene family (TprB, TprH, TprL, Tpp15) (Table 2) were selected (Weinstock et al. 1998). Long-range PCR products of these genes were produced from modern *T. pallidum pallidum* DNA and were subsequently used as bait for the molecular enrichment through hybridization following the Maricic et al. protocol for long-range PCR bait production (Table SI.2) (Maricic, Whitten, and Pääbo 2010). The following primers were designed: TprB-FWD: CGTATATGCGCGTTTTGCGT, TprH-RVS: ACCATGTAGGGCGTTGGTTC, TprH-FWD: GAGTACGCTGTCCCTGTCTT, TprH-RVS: ACCATGTAGGGCGTTGGTTC, TprL-FWD: CAAGGACCTGACCGTTGATT, TprL-RVS: CGCACACAAAGGTGCCAAA, Tpp15-FWD: ATGGTGAAAAGAGGTGGCG, Tpp15-RVS: CTACCTGCTAATAATGGCTTCCT.

PCRs reactions were carried out using the Roche long-range PCR kit in 100 μl reactions (1μl DNA template, 1 unit 10× PCR buffer 2, 0.4 mg/mL BSA, 3μl DMSO, 3125 μM each dNTP, 7U polymerase and 0.3 mM each primer) with the following thermal profile: an initial denaturation at 92°C for 2 min, 10 cycles consisting of a denaturation at 92°C for 10 sec, an annealing at 66°C for 30 sec and a 9-min elongation at 68°C, followed by 30 cycles using the same thermal profile with an additional increase of the elongation time by 20 sec each cycle and a final elongation at 68°C for 7 min. After purification using MinElute spin columns and quantification by NanoDrop, the PCR products were sonicated to 300 bp using a S220 Covaris (Duty cycle 10%, peak incident power 140W, cycles per burst 200, time 120 sec). Products were ligated to biotinylated adapters and immobilized on streptavidin-coated magnetic beads.

Between 8 to 10 individually indexed libraries were pooled at a final concentration of 2 µg of DNA. For the blocking step, a blocking oligos mix composed of 500 µM of each blocking oligo primer Bio4, Bio6, Bio8 and Bio10 (Maricic, Whitten, and Pääbo 2010). Negative controls (extraction and library blanks) were pooled and enriched separately. After 48 h incubation at 65°C and several washing steps, the library molecules were eluted by NaOH melting and quantified by qPCR (LightCycler 480, Roche) following previous methods (Maricic, Whitten, and Pääbo 2010). Prior to sequencing, sample pools and negative controls were combined equimolarly to a final concentration of 10nM. The pools were sequenced using a NextSeq 500 Illumina platform at the Functional Genomic Center of Zurich (Switzerland).

### **Supplementary Note 2 - Analysis**

#### **Metagenomic analysis**

The metagenomic composition was reconstructed using MALT, which uses the NCBI taxonomy to provide unique names and IDs for over 660,000 taxa, including approximately 25,000 prokaryotes. We executed MALT with the following mapping parameters: reads with a minimum 85% identity (−−minPercentIdentity) were considered as a possible match to the reference; only nodes with minimum support of five reads (−−minSupport set to 5); BlastN mode and SemiGlobal alignment were applied and a top percent value of 1 (−−topPercent). The remaining parameters were set to default  (Vågene et al. 2018). Results were visualized with MEGAN v. 11.7 (Huson et al. 2007). Sample comparisons used normalized counts to control for variation in sequencing depth. Further visualization and figures were performed with the R package phyloseq (McMurdie and Holmes 2013). Genera that were found in the blanks and that are known to be common residual contaminants in the reagents were not taken into account, mainly Pseudomonas and Burkholderia (Salter et al. 2014); one of the blanks had an abundance higher than 90% of the genus Plantactinospora, a very resistant spore-forming member of the Actinobacteria, but since it was absent in the other blanks we assumed it to be a punctual contamination.

The proportion of taxa present in each fixative sample that can be explained by tissue contribution or laboratory contamination was estimated by using SourceTracker2 (Knights et al. 2011). A BIOM1 file was generated from the MEGAN output, complemented with a mapping file where fixative samples were defined as ‘sink’ material and each tissue sample and laboratory blank as ‘source’. A sink rarefaction depth of 0 and source rarefaction depth of 0 were used to maximize taxa trace detection. The mixing proportions output was then plotted in a stacked bar chart graph using R (Core Team and Others 2013).

#### **Endogenous DNA - Methods**

To assess the levels of endogenous DNA in the samples, we used the EAGER pipeline  (Peltzer et al. 2016) to perform an alignment against both mitochondrial and whole human genome. An important characteristic of ancient sequencing libraries is the occurrence of the

substitution of C by T at the fragment ends (Briggs et al. 2007). This is due to the post-mortem decay of the DNA and can be used to authenticate ancient DNA. We selected the in-built function DamageProfiler in EAGER for an estimation of damage in 5’ and 3’ ends for ancient DNA authentication (Neukamm, Peltzer, and Nieselt 2020).  Reads were adapter clipped, merged (suited for pair-end datasets), and quality trimmed using Clip&Merge (Peltzer n.d.). For those read pairs that could not be merged because the overlap region was shorter than 10 nucleotides, or for which the corresponding read was removed during the combined adapter clipping and quality filtering step, the respective single-end reads were first trimmed at the 3’ end such that all bases have a Phred quality score of at least 20 and then mapped individually. The resulting reads for all samples were mapped to the human mitochondrial reference genome hg19 (NC_001807) treated as double end reads and using the CircularMapper feature built in the EAGER pipeline, with an error rate (-n) of 0.2 to assure high specificity (Peltzer n.d.).  The enrichment factor for all enriched libraries was calculated by dividing the percentage of endogenous DNA after enrichment by the percentage of endogenous DNA in the shotgun sequencing (Tables SI.7, SI.8, SI.9, SI.10).

#### **Pathogen DNA Capture - Methods**

To assess the presence of pathogen DNA in the samples, we used the EAGER pipeline  (Peltzer et al. 2016) to perform an BWA alignment against the whole genome of *Mycobacterium leprae*, *Mycobacterium tuberculosis*, and *Treponema pallidum.* We selected the built-in function DamageProfiler in EAGER for an estimation of damage in 5’ and 3’ ends for ancient DNA authentication (Neukamm, Peltzer, and Nieselt 2020).  Reads were adapter clipped, merged (suited for pair-end datasets), and quality trimmed using Clip&Merge (Peltzer n.d.). For those read pairs that could not be merged because the overlap region was shorter than 10 nucleotides or for which the corresponding read was removed during the combined adapter clipping and quality filtering step, the respective single-end reads were first trimmed at the 3’ end such that all bases have a Phred quality score of at least 20 and then mapped individually. Duplicate reads were merged using DeDup. The resulting reads for all samples were mapped treated as double end reads and using the CircularMapper and BWA features built in the EAGER pipeline, with an error rate (-n) of 0.2 to assure high specificity (Peltzer n.d.). The enrichment factor for all enriched libraries was calculated by dividing the percentage of endogenous DNA after enrichment by the percentage of endogenous DNA in the shotgun sequencing.

### **Supplementary Note 3 - Results**

#### **Endogenous DNA - Results**

For the shotgun mtDNA screening, nine out of fourteen tissue replicates showed detectable traces of mtDNA, eight with a 5X coverage above 98% and one above 24% (Table SI.2). These replicates demonstrate the characteristic misincorporation patterns for ancient DNA (elevated C to T substitutions at 5′-ends and elevated G to A substitutions at 3′-ends) in frequencies ranging from 10- 20% damage (Neukamm, Peltzer, and Nieselt 2020). For the liquid samples, only sample HA4.1 showed sufficient DNA to be considered as ancient (94.59% 5X coverage of mitogenome, 16% damage). The enrichment strategy (Maricic, Whitten, and Pääbo 2010) (SI, Notes 2) was powerful enough to increase the endogenous content of all the tissue samples (Table SI.3), with fold increments from 14 to 275, and also for eight of the ten liquid samples (Table SI.4), not including HA4.1. This sample had a remarkable increase with a final output of more than three million mapped ancient mtDNA reads, a 537-fold increase in the endogenous content (100% coverage) and retaining a damage of 10%. Seven tissue samples and liquid sample HA4.1 were assigned to human mtDNA haplogroups, and the percentage of contaminated DNA was estimated (Table SI.5). The haplogroups identified were typically European (Table SI.2) (Renaud et al. 2015; Weissensteiner et al. 2016).

#### **Pathogen DNA Capture - Results**

For the tuberculosis samples, the tissue P1255 was the most successful with 1287 reads that corresponded to 0.25% endogenous content, and presented a damage pattern of 10%, although with a quite low coverage (0.92%) and a high cluster factor (2.34). The fixative sample for P1255, however, provided a very low number of reads - only 151 with a 2.7% endogenous content, 0.04 coverage and cluster factor of 6.34, that were not enough to be discerned as ancient or not. For the leprosy samples, the tissue HA4.1 delivered the highest output with 1051 reads with a damage of 16%, (0.27% endogenous content, 0.93 coverage) but with an extremely high cluster factor of 25.11, indicating an elevated number of repetitive reads. For the fixative material from HA4.1, only 790 reads were detected, with a damage pattern of 22% (0.082% endogenous DNA, 0.3 coverage, and a cluster factor of 10.29), not providing enough material to allow for a finer analysis of the pathogen. An interesting finding was sample P1245 that presented the second-best results for the leprosy enrichment with 917 reads, 20% damage pattern (0.34% endogenous content, 2.4 coverage and a cluster factor of 13), although it was supposedly diagnosed with syphilis (Table SI.1).

### **Supplementary Note 4 - Extended Sample Information**

#### **Notes on the History of HA4.1 (RCS Reference number: RCSPC/03240)**

by Martyn Cooke

Specimen HA 4.1 was donated to the RCS in 1886 and consists of the tongue and adjacent tissues of a man suffering from leprosy. The patient history indicates that the man died in the Trinidad Leper Asylum, so we must assume that the specimen was removed at post-mortem and placed in a tissue fixative of some sort of alcohol or ‘spirit’ in Trinidad. Spirit solutions had been in regular usage since the 1600’s and most anatomical or medical museums at that time would have been using a solution of Ethyl Alcohol. However, we can only guess at what was available in The Trinidad Leper Asylum to preserve specimen HA 4.1. It was clearly a suitable substance, as it clearly minimized autolytic changes and decomposition during the long journey to the UK.

Transfer to Kaiserling III Preservative

At the beginning of the 1900’s, the Kaiserling method was one of many methods to replace Spirit preparations and become a standard method for many pathology museums throughout the world, based around the use of Formaldehyde as a direct replacement to Alcohol. Formaldehyde was first prepared in 1859 and by the time Kaiserling’s new method was introduced in 1896 (Kaiserling 1896), Formaldehyde was already going into commercial production. Kaiserling made further changes to his method in 1897 (Kaiserling 1897) and again in 1900 (Kaiserling 1900). Formaldehyde fixation does initially cause a loss of colour due to oxidation, but this process can be partially reversed.

Kaiserling’s three stage procedure is as follows:

- **Kaiserling I (KI) solution:** The Fixative. Formaldehyde and buffering salts.
- **Kaiserling II (KII) solution:** Colour Restoration. Short immersion in Alcohol.
- **Kaiserling III (KIII) solution:** The Preservative. Glycerine and buffering salts.

At the same time Kaiserling was perfecting his method, many other scientists were developing their own solutions and modifications along very similar, if not identical lines. It is extremely difficult to determine the exact makeup and chemical ingredients of KIII preservatives that have been used over time. It is also very difficult to determine when specimen HA4.1 was transferred from Alcohol to KIII. Without any records to refer to, it is almost impossible to determine what happened to the specimen. Some records related to the purchase of Glycerine for specimen preparation and the earliest dated back to 1941, so we can say with a reasonable amount of accuracy that the RCS had the means to prepare and use Glycerine in Kaiserling III Preservative as early as the 1940’s.

In 1951 the RCS was actively using and promoting a variation of the Kaiserling technique and L. Proger, the Pathological Curator of the RCS produced a short article (Annals of the Royal College of Surgeons of England Volume 8, No 5, May 1951, pages 388-391). The chemical formula of the preservative (Described as the ‘Mounting Solution’) is as follows:

The Pick-Judah Method (Daukes’ Modification)

- Sodium Acetate 2.7 Kg
- Distilled Water 9 L
- Glycerine 5.4	L
- Camphor 50g dissolved in 200 mL Industrial Methylated Spirit.

This raises the concern that the ‘Kaiserling III’ preservative possibly used for preservation of specimen HA 4.1 may not even have been Kaiserling’s own method, but an inspired variant.

In 1988, the RCS introduced a number of modifications to the Kaiserling fixation/preservation process:

- The original Kaiserling I formaldehyde fixative was replaced by a histological formaldehyde fixative, containing alternate and improved buffering salts.
- The original Kaiserling II colour restoration in an alcohol bath was replaced by the addition of chemical (Reducing) agents into the preservative to restore colour
- The original Kaiserling III preservative was altered to give a higher percentage of Glycerine, which reduced the likelihood of mould growth and eliminated the need for Camphor or Formaldehyde additives.

This means that the original Kaiserling III preservative formula that was used in the 1950’s for specimen HA4.1 was not the same Kaiserling III preservative formula that was used in the 1990’s.

Previous Research Request for Sampling HA 4.1

During the 1990’s, a team approached the RCS from St Mary’s Hospital, who were investigating new methods of obtaining DNA from *Mycobacterium* and requested a sample from specimen HA 4.1. When sampled, it was found that the sample was mounted in pure Liquid paraffin. Liquid Paraffin is a highly refined mineral oil derived from petroleum. It is also known as Medicinal Paraffin (due to its more historical use as a laxative), Paraffinum Liquidum and Russian Mineral Oil. Liquid Paraffin is a colourless, odourless, inert substance, which has found favour as a tissue preservative in certain circumstances.

The RCS was possibly unique in using Liquid Paraffin almost exclusively for routine pathological specimens for a period of time, as it minimized discoloration of the preservative caused by pigmentation leaching out of the specimen. Histological examination showed Liquid Paraffin to be a very good preservative. However, the high refractive index had a tendency to cause muscle and skin to darken and become slightly transparent, which obscured many pathological lesions. For this reason, many paraffin preserved specimens were returned back to Kaiserling III preservative, where the transparency was reversed.

Liquid Paraffin was used from 1978 onwards and started to be phased out for routine use by the 1990’s. It is possible that Specimen HA 4.1 was transferred from Kaiserling III preservative to Liquid Paraffin around 1978 and following the first sampling session in the 1990’s was returned back to Kaiserling III preservative, where it remained until the current research study sampling took place.

Brief Summary of HA4.1 fixative history

The current RCS Kaiserling III preservative formulae is as follows:

- Distilled Water 15,000mL
- Glycerine 10,000mL
- Sodium Acetate Trihydrate 3,500g

The timeline of fluid changes suggested is as follows:

- 1886 -	Fixation, in an unknown Alcohol solution in Trinidad.
- 1886 -	Preservation, transfer to 70-80% Ethyl Alcohol at RCS
- 1950’s - Preservation, transfer to Pick, Judah, Daukes preservative (Kaiserling III Modification)
- 1970’s - Preservation, transfer to pure Liquid Paraffin
- 1990’s - Preservation, transfer to Current RCS Kaiserling III preservative to the present time (2021)

### **Supplementary Tables**

**Table SI.1:** List of tissue and fixative samples utilized in this study.

* Based on the jar lid sealing, these samples have not undergone recent conservation, i.e. the fixative was not recently replaced.

| **Sample** | **Diagnosis** | **Tissue** | **Fixative** | **Dating** |
| --- | --- | --- | --- | --- |
| HA4.1 | Leprosy | Lymph node | Kaiserling | 1886 |
| R15.3 | Syphilis | Larynx cartilage | 70%EtOH/silicone | 1877 |
| P906 | Tuberculosis | Femur | 70%EtOH/silicone | 1760-1793 |
| P1189 | Syphilis | Lung | 70%EtOH * | 1760-1793 |
| P1245 | Syphilis | Larynx | 70%EtOH * | 1760-1793 |
| P1255 | Tuberculosis | Lung | 70%EtOH * | 1760-1793 |
| P1499 | Molluscum | Eyelid skin | 70%EtOH * | 1760-1793 |
| P1505 | Wart | Genital wart | 70%EtOH/silicone | 1760-1793 |
| P1526 | Carcinoma | Lip carcinoma | 70%EtOH * | 1760-1793 |
| S51.10 | Syphilis | Femur head | 70%EtOH * | 1889 |

**Table SI.2:** Primer list: Primers used for the bait design of the *Treponema pallidum* hybridization-capture protocol.

| **Primer** | **Sequence** | **Fragment length** |
| --- | --- | --- |
| TprB fwd | CGTATATGCGCGTTTTGCGT | 1979 bp |
| TprB rev | TCACCACAGAACCTTACACGA | 1979 bp |
| TprH fwd | GAGTACGCTGTCCCTGTCTT | 2078bp |
| TprH rev | ACCATGTAGGGCGTTGGTTC | 2078bp |
| TprL fwd | CAAGGACCTGACCGTTGATT | 1541bp |
| TprL rev | CGCACACAAAGGTGCCAAA | 1541bp |
| Tpp15 fwd | ATGGTGAAAAGAGGTGGCG | 425bp |
| Tpp15 rev | CTACCTGCTAATAATGGCTTCCT | 425bp |

*Fwd: forward; Rev: reverse

**Table SI.3:** Long-range PCR reaction for the generation of baits.

| **LR-PCR** | | | | | |
| --- | --- | --- | --- | --- | --- |
| **Kit Nr** | **Reagent** | **Concentration** | **Final** | **ul** | **4.4** |
| #2 (on ice) | Buffer 2 green | 5 x | 1x | 20 | 88 |
| #5 | DMSO | 100% | 3% | 3 | 13.2 |
| #6 (on ice) | dNTP mix | 10uM | 0,5ulM | 5 | 22 |
|  | BSA | 10mg/ml | 0,4mg/ml | 4 | 17.6 |
|  | Primer For | 10uM | 0,3uM | 3 | 13.2 |
|  | Primer Rev | 10uM | 0,3uM | 3 | 13.2 |
| #1 (on ice) | Enzyme mix | 5U/ul | 0,075U | 1.4 | 6.16 |
|  | H20 |  |  | 59.6 | 262.24 |
|  | Template ( Modern syphilis DNA) |  |  | 1 | 4.4 |
|  | Total |  |  | 100 | 440 |

**Table SI.4:** Temperature settings for the LR-PCR

| **Temperature profile** | | |
| --- | --- | --- |
| Temp | Time | Cycles |
| 92°C | 2min |  |
| 92°C | 10sec | 10 cycles |
| 66°C | 30sec |  |
| 68°C | 9min |  |
| 92°C | 10sec | 24 cycles |
| 66°C | 30sec |  |
| 68°C | 9min ( +20 sec/cycle) |  |
| 68°C | 7min |  |
| 8°C | Hold |  |

**Table SI.5:** Shotgun screening for mtDNA, indicating the cluster factor and the percentage of coverage (1X and 5X).

| **Sample** | **Cluster Factor** | **Coverage >= 1X in %** | **Coverage >= 5X in %** |
| --- | --- | --- | --- |
| HA4.1 (L) | 1.104 | 100 | 94.59 |
| HA4.1 (LN) | 1.194 | 31.12 | 0 |
| HA4.1 (T) | 1.194 | 99.99 | 98.39 |
| P1189 (L) | 3 | 0.59 | 0 |
| P1189 (T) | 1.758 | 100 | 99.6 |
| P1245 (L) | 6.167 | 1.79 | 0 |
| P1245 (T) | 1.539 | 100 | 100 |
| P1255 (T) | 1.102 | 100 | 99.97 |
| P1499 (L) | NA | NA | NA |
| P1499 (T) | 1 | 30.56 | 0 |
| P1505 (L) | NA | NA | NA |
| P1505 (T) | 1.257 | 76.52 | 24.87 |
| P1526 (L) | NA | NA | NA |
| P1526 (T) | 2.007 | 100 | 99.99 |
| P906 (L) | NA | NA | NA |
| P906 (B) | 1 | 0.87 | 0 |
| P906 (T) | 2.525 | 18.99 | 0 |
| R15.3 (L) | NA | NA | NA |
| R15.3 (C) | 1.12 | 100 | 100 |
| R15.3 (CS) | 1.233 | 100 | 100 |
| R15.3 (M) | NA | NA | NA |
| S51.10 (L) | 2 | 0.91 | 0 |
| S51.10 (T) | 1.098 | 100 | 99.89 |

L= Liquid; T = Tissue; LN = Lymph Node; B = Bone; C = Cartilage; CS = Cross Section; M= Mucosa

**Table SI.6:** Endogenous content of the mtDNA present in the tissue samples before and after enrichment, and the calculated fold increase.

| **Sample** | **Shotgun Endogenous DNA (%)** | **Enriched Endogenous DNA (%)** | **Fold increase** |
| --- | --- | --- | --- |
| HA.4.1LN | 0.006 | 0.207 | 34.5 |
| HA4.1T | 0.09 | 7.223 | 80.26 |
| P1189T | 0.22 | 41.593 | 198.06 |
| P1245T | 1.251 | 47.769 | 38.19 |
| P1255T | 0.135 | 28.512 | 211.2 |
| P1499T | 0.002 | 0.253 | 126.5 |
| P1505T | 0.021 | 4.884 | 232.57 |
| P1526T | 0.456 | 43.575 | 95.56 |
| P906B | 0.001 | 0.014 | 14 |
| P906T | 0.003 | 0.531 | 177 |
| R15.3CS | 0.839 | 25.064 | 29.9 |
| R15.3LN | 0.66 | 22.244 | 33.7 |
| S51.10T | 0.092 | 25.361 | 275.7 |

L= Liquid; T = Tissue; LN = Lymph Node; B = Bone; C = Cartilage; CS = Cross Section; M= Mucosa

**Table SI.7:** Endogenous content of the mtDNA present in the fixative samples before and after enrichment, and the calculated fold increase.

| **Sample** | **Shotgun Endogenous DNA (%)** | **Enriched Endogenous DNA (%)** | **Fold increase** |
| --- | --- | --- | --- |
| HA4.1 | 0.083 | 44.612 | 537.5 |
| P1189 | 0.000 | 1.621 | - |
| P1245 | 0.000 | 2.249 | - |
| P1255 | 0.002 | 18.639 | 9319.5 |
| P1499 | 0.000 | 0.000 | - |
| P1505 | 0.000 | 0.751 | - |
| P1526 | 0.000 | 1.987 | - |
| P906 | 0.000 | 0.732 | - |
| R15.3 | 0.000 | 0.000 | - |
| S51.10 | 0.000 | 1 | - |

**Table SI.8:** Mitochondrial DNA haplogroup estimation after the hybridization capture, indicating quality of the estimate and percentage of the contamination, and percentage of damage in the reads.

| **Sample** | **Haplogroup** | **Quality (%)** | **Cont.Est** | **Damage (%)** |
| --- | --- | --- | --- | --- |
| HA4.1 (T) | H2a2 | 52.20 | 0.01 | 23 |
| HA4.1 (LN) | H2a2 | 51.69 | 0.01 (?) | failed |
| HA4.1 (L) | H | 91.44 | 0.04 (K1) | 10 |
| P906 (T.1) | K1 | 66.67 | 0.99 (?) | 9 |
| P906 (T.2) | failed | failed | failed | failed |
| P906 (L) | failed | failed | failed | failed |
| P1189 (T) | I2 | 94.77 | 0.01 (?) | 17 |
| P1189 (L) | failed | failed | failed | failed |
| P1245 (T) | U5a1a1e | 98.81 | 0.02 (?) | 23 |
| P1245 (L) | failed | failed | failed | failed |
| P1255 (T) | H1bb | 97.54 | 0.00 | failed |
| P1255 (L) | failed | failed | failed | failed |
| P1499 (T) | H1bb | 65.06 | 0.01 (?) | 20 |
| P1499 (L) | failed | failed | failed | failed |
| P1505 (T) | W3a1a2 | 76.55 | 0.01 (?) | 21 |
| P1505 (L) | failed | failed | failed | failed |
| P1526 (T) | H3 | 94.57 | 0.01 (?) | failed |
| P1256 (L) | failed | failed | failed | failed |
| R153 (C) | K1e1 | 94.19 | 0.00 | failed |
| R153 (L) | failed | failed | failed | failed |
| S51.10 (T) | failed | failed | failed | failed |
| S51.10 (L) | failed | failed | failed | failed |

L= Liquid; T = Tissue; LN = Lymph Node; B = Bone; C = Cartilage; (?) = unknown contaminant

**Table SI.9:** Endogenous content of the pathogen DNA present in the tissue samples before and after enrichment, and the calculated fold increase.

| **Sample** | **Shotgun DNA (%)** | **Enriched DNA (%)** | **Fold increase** |
| --- | --- | --- | --- |
| HA4.1LN (leprosy) | 0.003 | 0.008 | 2.66 |
| HA4.1T (leprosy) | 0.011 | 0.273 | 24.81 |
| P1189T (syphilis) | 0.003 | 0.005 | 1.666 |
| P1245T (syphilis) | 0.001 | 0.002 | 2 |
| R15.3CS (syphilis) | 0.000 | 0.000 | - |
| R15.3LN (syphilis) | 0.000 | 0.000 | - |
| S51.10T (syphilis) | 0.000 | 0.000 | - |
| P1255T (TB) | 0.013 | 0.248 | 19.076 |
| P906B (TB) | 0.004 | 0.004 | 2 |
| P906T (TB) | 0.006 | 0.017 | 2.83 |

**Table SI.10:** Endogenous content of the pathogen DNA present in the fixative samples before and after enrichment, and the calculated fold increase.

| **Sample** | **Shotgun DNA (%)** | **Enriched DNA (%)** | **Fold increase** |
| --- | --- | --- | --- |
| HA4.1 (leprosy) | 0.013 | 0.082 | 6.30 |
| P1189 (syphilis) | 0.002 | 0.006 | 3 |
| P1245 (syphilis) | 0.003 | 0.005 | 1.66 |
| R15.3 (syphilis) | 0.001 | 0.006 | 6 |
| S51.10 (syphilis) | 0.003 | 0.005 | 1.66 |
| P1255 (TB) | 0.019 | 0.027 | 1.41 |
| P906 (TB | 0.007 | 0.0352 | 5.02 |

### **Supplementary Figures**

**Figure SI.1:** Richness of bacteria expressed with alpha diversity indices observed (top) and Shannon Index (bottom), both separated based on type of fixative (EtOH70 in green, Kaiserling in blue) and negative controls (Blank in red).

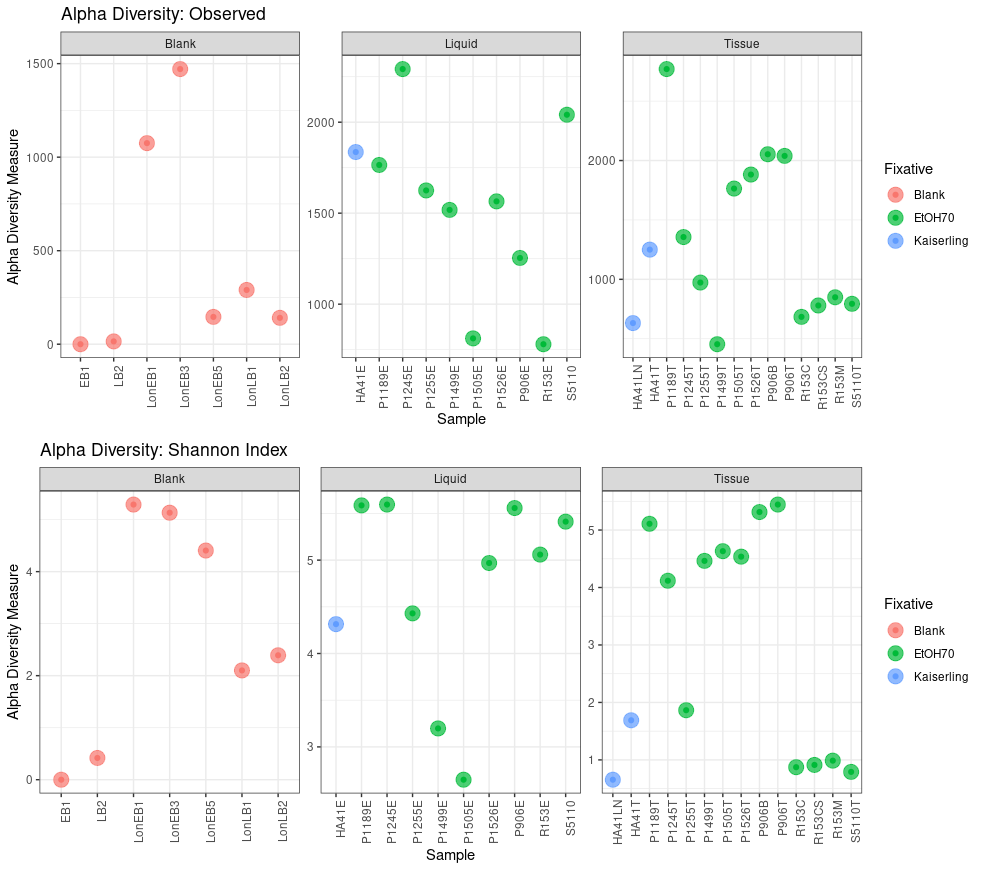
EB = Extraction Blank; LB = Library Blank; Lon = London; E = Ethanol (liquid) ; LN = Lymph Node; T = Tissue; B = Bone; C = Cartilage; CS = Cross Section; M = Mucosa.

**Figure SI.2:** Main 20 bacterial families present in the samples expressed in their percentage of abundance, separated by blank, liquid or tissue samples.
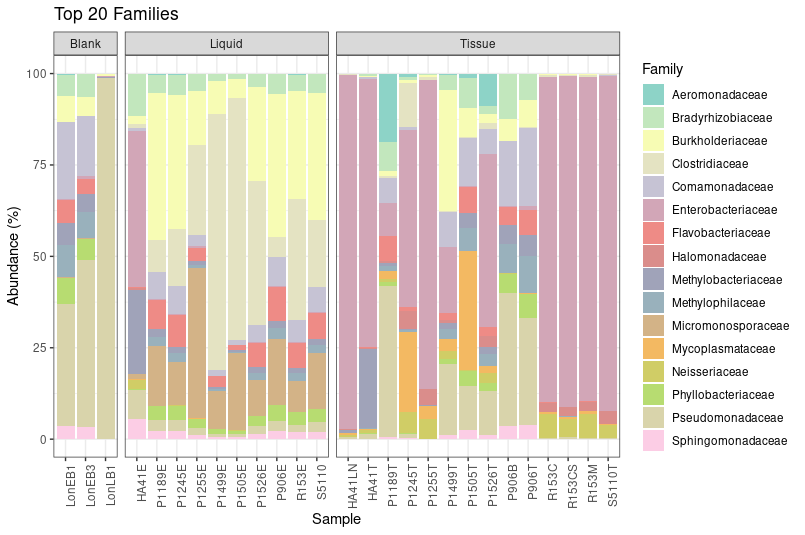
EB = Extraction Blank; LB = Library Blank; Lon = London; E = Ethanol; LN = Lymph Node; T = Tissue; B = Bone; C = Cartilage; CS = Cross Section; M = Mucosa.

Kaiserling, Carl. 1896. “Über Die Conservierung von Sammlungspräpaten Mit Erhaltung Der Natürlichen Farben.” *Berliner Klinische Wochenschift* 33 ((35)): 775–77.

———. 1897. “Weitere Mittheilungen über Die Herstellung Möglichst Naturgefreuer Sammlungspräpaten.” *Virchows Archiv Fur Pathologische Anatomie Und Physiologie Und Fur Klinische Medizin* 147 ((3)): 389–417.

———. 1900. “Über Die Konservierung Und Aufstellung Pathologisch-Anatomischer Präparate Für Schau- Und Lehrsammlungen.” In *Verhandlungen Der Deutschen Pathologischen Boyle.* Baltimore: John Hopkins University Press.

Kircher, Martin, Susanna Sawyer, and Matthias Meyer. 2012. “Double Indexing Overcomes Inaccuracies in Multiplex Sequencing on the Illumina Platform.” *Nucleic Acids Research* 40 (1): e3.

Knapp, Michael, and Michael Hofreiter. 2010. “Next Generation Sequencing of Ancient DNA: Requirements, Strategies and Perspectives.” *Genes* 1 (2): 227–43.

Knights, Dan, Justin Kuczynski, Emily S. Charlson, Jesse Zaneveld, Michael C. Mozer, Ronald G. Collman, Frederic D. Bushman, Rob Knight, and Scott T. Kelley. 2011. “Bayesian Community-Wide Culture-Independent Microbial Source Tracking.” *Nature Methods* 8 (9): 761–63.

Llamas, Bastien, Guido Valverde, Lars Fehren-Schmitz, Laura S. Weyrich, Alan Cooper, and Wolfgang Haak. 2017. “From the Field to the Laboratory: Controlling DNA Contamination in Human Ancient DNA Research in the High-Throughput Sequencing Era.” *STAR: Science & Technology of Archaeological Research* 3 (1): 1–14.

Maricic, Tomislav, Mark Whitten, and Svante Pääbo. 2010. “Multiplexed DNA Sequence Capture of Mitochondrial Genomes Using PCR Products.” *PloS One* 5 (11): e14004.

McMurdie, Paul J., and Susan Holmes. 2013. “Phyloseq: An R Package for Reproducible Interactive Analysis and Graphics of Microbiome Census Data.” *PloS One* 8 (4): e61217.

Meyer, Matthias, and Martin Kircher. 2010. “Illumina Sequencing Library Preparation for Highly Multiplexed Target Capture and Sequencing.” *Cold Spring Harbor Protocols* 2010 (6): db.prot5448.

Neukamm, Judith, Alexander Peltzer, and Kay Nieselt. 2020. “DamageProfiler: Fast Damage Pattern Calculation for Ancient DNA.” *Cold Spring Harbor Laboratory*. https://doi.org/[10.1101/2020.10.01.322206](http://dx.doi.org/10.1101/2020.10.01.322206).

Peltzer, Alexander. n.d. *CircularMapper*. Github. Accessed January 26, 2021a. <https://github.com/apeltzer/CircularMapper>.

———. n.d. *ClipAndMerge*. Github. Accessed January 26, 2021b. <https://github.com/apeltzer/ClipAndMerge>.

Peltzer, Alexander, Günter Jäger, Alexander Herbig, Alexander Seitz, Christian Kniep, Johannes Krause, and Kay Nieselt. 2016. “EAGER: Efficient Ancient Genome Reconstruction.” *Genome Biology* 17 (March): 60.

Renaud, Gabriel, Viviane Slon, Ana T. Duggan, and Janet Kelso. 2015. “Schmutzi: Estimation of Contamination and Endogenous Mitochondrial Consensus Calling for Ancient DNA.” *Genome Biology* 16 (October): 224.

Schuenemann, Verena J., Pushpendra Singh, Thomas A. Mendum, Ben Krause-Kyora, Günter Jäger, Kirsten I. Bos, Alexander Herbig, et al. 2013. “Genome-Wide Comparison of Medieval and Modern Mycobacterium Leprae.” *Science* 341 (6142): 179–83.

Vågene, Åshild J., Alexander Herbig, Michael G. Campana, Nelly M. Robles García, Christina Warinner, Susanna Sabin, Maria A. Spyrou, et al. 2018. “Salmonella Enterica Genomes from Victims of a Major Sixteenth-Century Epidemic in Mexico.” *Nature Ecology & Evolution* 2 (3): 520–28.

Weinstock, G. M., J. M. Hardham, M. P. McLeod, E. J. Sodergren, and S. J. Norris. 1998. “The Genome of Treponema Pallidum: New Light on the Agent of Syphilis.” *FEMS Microbiology Reviews* 22 (4): 323–32.

Weissensteiner, Hansi, Dominic Pacher, Anita Kloss-Brandstätter, Lukas Forer, Günther Specht, Hans-Jürgen Bandelt, Florian Kronenberg, Antonio Salas, and Sebastian Schönherr. 2016. “HaploGrep 2: Mitochondrial Haplogroup Classification in the Era of High-Throughput Sequencing.” *Nucleic Acids Research* 44 (W1): W58–63.
